# MicroRNA-224 Promotes Cell Migration and Invasion by Targeting HOXA5 expression in Hepatocellular Carcinoma

**DOI:** 10.1101/2020.08.27.269654

**Authors:** Yuwu Liu, Chen Lin, Dongmei Wang, Sailan Wen, Junpu Wang, Zhihong Chen, Jing Du, Chao Ning, Deyun Feng

## Abstract

MicroRNAs (miRNAs) are involved in the regulation of multiple cellular pathways and play a key role in the development and progression of tumor. Based on the cellular function of their targets, miRNAs play the role of oncogenes or tumor suppressor genes. Multiple studies have shown that abnormal expression of miRNAs has close relation with the incidence of HCC, but the mechanism of miRNAs in HCC still needs further research. In the present study, we showed that the overexpression of miR-224 can reduce the mRNA and protein expression of ADAM17 and HOXA5, the silencing of miR-224 can increase the protein expression of ADAM17 and HOXA5. Dual luciferase reporter assays showed that miR-224 can directly regulate the expression of ADAM17 and HOXA5. Importantly, we found that miR-224 positively regulates cell migration and invasion in HCC, miR-224 overexpression can promote the migration and invasion of BEL-7402 cell, and miR-224 silencing can suppress the migration and invasion of BEL-7402 cell. miR-224 overexpression can result in the redistribution of cell cycle, the cell percentage of S phase was increased significantly, the cell percentage of G1 phase was decreased significantly, and there is no noticeable change for the cell percentage of G2 phase. These results demonstrated that it may be exert the function of oncogenes in a particular link of cancer cell growth. In conclusion, these results suggest that miR-224 will become a promising biological target in the treatment strategy of hepatocellular carcinoma (HCC).

## Introduction

MicroRNAs(miRNAs), a member of the non-coding RNA family, are 19 to 25 nt-sized transcripts cleated from 70-100 nucleotide (nt) -sized hairpin shaped precursors. Among distantly related organisms, the sequence of miRNAs is conserved, suggesting that these molecules are involved in the basic biological processes of the organism [1-3]. To date, more than 500 miRNAs have been predicted to be expressed in humans [4, 5], and these miRNAs are estimated to regulate the expression of over 5,000 human genes or 30% of human proteins[6]. Interactions between MiRNAs and their numerous target genes may represent important levels of gene regulation in cells[7]. Although the exact function of miRNAs is still unknown, existing studies have shown that miRNAs have a variety of different expression patterns and can modulate many different developmental and physiological processes. In addition, dysregulation of miRNAs expression may lead to human disease[8]. Several studies have shown that close to 50% of miRNAs genes are located in cancer-related genomic regions or fragile sites, which draws attention to the importance of miRNAs in cancer [9-11].

Abnormal expression of MiRNAs has been shown to involve a variety of solid tumors, including hepatocellular carcinoma, colon cancer, prostate cancer, breast cancer, ovarian cancer and lung cancer[12-15]. In addition, abnormal expression of miRNAs has been found to be associated with postoperative survival of lung cancer patients[16], and can be used as a biomarker for the diagnosis and treatment of lung cancer[17]. According to the role of their target genes, miRNAs can play an important role in the occurrence and development of tumors as both tumor suppressor genes and oncogenes[18]. The use of gene chip make it possible for the analysis of large-scale miRNAs expression in tumor study, we can confirm a lot of miRNAs in cancer tissues and normal tissues in the differences in expression by this way, so that we can better explain the imbalance in the development of tumor miRNAs. Although the function of some cancer-related miRNAs has been revealed, the mechanism of action of most miRNAs in tumors remains unclear[19].

Some miRNAs have been proved to have the function of tumor suppressor genes. Johnson SM et al[20] reported that the let family can regulate the expression of RAS gene, revealing that it may play an inhibitory role in the development of lung cancer. O ‘donnell et al[21] reported that miR-17-5p and miR-20a negatively regulate the expression of E2F1, thereby affecting the function of c-myc gene and ultimately controlling the signals of proliferation. Some miRNAs have been shown to have the function of oncogenes, and miR-21 has been found to be up-regulated in both glioma and breast cancer and to play an anti-apoptotic role[22, 23]. MiR-372 and miR-373 play the role of oncogenes in human testicular germ cell tumor, and in the presence of wild-type P53, they can inhibit the P53 signaling pathway and allow the proliferation of tumor cells[24]. Multiple studies have shown that miR-224 is overexpressed in human tumors, such as colon cancer [25, 26], uterine cancer[27], glioma[28], breast cancer[29] and hepatocellular carcinoma[30]. This suggests that up-regulated miR-224 may play an important role in tumor formation. miR-224 can affect several key physiological functions of cancer cells, including apoptosis[31], proliferation, migration and invasion[32, 33], which proves its potential carcinogenicity.

Although we have some understanding of the expression and function of miR-224 in hepatocellular carcinoma based on some previous reports, the exact mechanism or intracellular target of miR-224 in hepatocellular carcinoma is still poorly understood. Therefore, the detection of the target of miR-224 is very important for our in-depth understanding of the mechanism of action of miR-224. In this study, we will further investigate the mechanism of miR-224 in the development of hepatocellular carcinoma. We screened the target genes of miR-224 through bioinformatics, and selected ADAM17 and HOXA5 as the starting points of this study, hoping to find strong evidence that miR-224 plays an important role in the development of hepatocellular carcinoma.

## Materials and Methods

### Patient samples

Paired hepatocellular carcinoma (HCC) tissues and the corresponding nearby noncancerous livers were obtained from 19 patients who underwent liver resection. Tissue samples were immediately snap-frozen in liquid nitrogen, and long-term stored at minus 80 degrees Celsius in refrigerator. These 19 samples were mainly used for fluorescence quantitative PCR detection. In addition,In addition, the paraffin-embedded samples of 42 cases of hepatocellular carcinoma and 8 cases of hepatic hemangioma were collected from the specimen bank archived in the Department of Pathology, Xiangya Hospital, Central South University. The normal liver tissue incidental to a specimen of hepatic hemangioma were used as control of HCC and adjacent nontumorous tissues. The 42 samples of hepatocellular carcinoma and 8 samples of hepatic hemangioma were mainly used for immunohistochemical detection.The written informed consents were provided for all of the patients, and all aspects of this study were approved by the Ethics Committee from Xiangya Hospital, Central South University, Changsha, China.

The clinicopathologic features of 42 patients with paraffin-embedded hepatocellular carcinoma samples were described briefly (S1 file). All of 42 patients contained 37 men and 5 women, the sex ratio(male/female) was 7:1. The mean age was 50 yeas old. According to microscopic pathological characteristics, histological grade of tumor differentiation was assigned: 17% were well-differentiated, 57% were moderately-differentiated, 26% were poorly-differentiated. Most of patients (36 cases) were single tumor, and only 5 cases were multiple tumors among 42 patients. The biggest size of tumor can be up to 20cm, the smallest was just 1cm. About 57% of the patients presented microscopic capsule and/or vascular invasion. Hepatic cirrhosis was recorded for 67% of the patients. Here we may draw a conclusion that cirrhotic liver is closely related to the occurrence of hepatocellular carcinoma.

### Cell culture

SMMC-7721, BEL-7402, Huh-7, HL-7702 (L-02) and HEK-293T were purchased from the Cell Bank of Chinese Academy of Sciences. SMMC-7721, BEL-7402 and HL-7702 (L-02) were cultured in RPMI 1640 medium (Gibco, Invitrogen) supplemented with 10% (v/v) fetal bovine serum (Gibco, Invitrogen), HuH-7 and HEK-293T were maintained in DMEM high glucose medium (Gibco, Invitrogen) supplemented with 10%(v/v) fetal bovine serum (Gibco, Invitrogen). All media contained 100 units of penicillin/ml and 100μg of streptomycin/ml. Cells were incubated at 37°C and supplemented with 5% CO2 in the humidified chamber.

### RNA oligonucleotides and transfection

miR-224 mimics and mimics Negative Control (mimics NC) were chemically synthesized by Shanghai GenePharma Co.Ltd (Shanghai, China). miR-224 inhibitor and inhibitor Negative Control(inhibitor NC) were obtained from Ambion, Inc (Austin, TX, USA). Cells transfection was performed using siPORT™ NeoFX™ Transfection Agent (ambition) following the manufacturer’s protocol.

### Real-time quantitative PCR

Total RNA was extracted from tissues and cultured cells using TRIzol (Invitrogen). Real-time quantitative PCR (qPCR) was carried out to quantify the expression of miR-224. TaqMan MicroRNA Transcription kit, TaqMan Universal Master Mix II and TaqMan asssys were used according to the manufacturer’s protocol (Applied Biosystems, USA). U6 snRNA was used as normalization control. Real-time quantitative PCR (qPCR) was performed to detect the expression of ADAM17 and HOXA5. Reverse transcription was carried out using the PrimeScript® RT reagent Kit with gDNA Eraser (TaKaRa), and cDNA products were amplified using SYBR ® Premix ExTaqTM II Kit (TaKaRa), GAPDH was served as an internal control for total cDNA content. The sequence of oligonucleotides used as PCR primers were: ADAM17 (Forward): 5’-GCT GAC CCA GAT CCC ATG AAG-3’, (Reverse): 5’-CTG CAT TAT CCC ATG AAG TGT TCC-3’. HOXA5(Forward): 5’-GAT GCG CAA GCT GCA CAT AAG-3’, (Reverse) : 5’-CGG GTC AGG TAA CGG TTG AA-3’. GAPDH(Forward) : 5’-GCA CCG TCA AGG CTG AGA AC-3’, (Reverse) : 5’-TGG TGA AGA CGC CAG TGG A-3’. The relative miRNA(miR-224) or mRNA(ADAM17 AND HOXA5) levels were calculated using the comparative cycle threshold [Ct, 2^-ΔΔCt^] method with their endogenous control.

### Immunohistochemistry (IHC)

We carried out the Immunohistochemistry of Paraffin-embedded sections according to a two-step protocol (Pllink-2 plus@ Polymer HRP Detection System, GBI). Briefly, the sections were first deparaffinized in xylene, then hydrated in a graded series of ethanol (100%∼50%) and tap water. The high pressure method was chosen to perform antigen retrieval in citrate buffer (0.01M, pH 6.0), subsequently the sections were incubated in 3% H2O2 at RT for 10 minutes so that endogenous peroxidase activity was blocked. After the sections were washed in PBS, treated with anti-ADAM17 antibody (Abcam, USA) at 4°C overnight. The sections were incubated with polymer helper at 37°C for 20 minutes, subsequently incubated with poly-HRP anti-Mouse IgG at the same temperature and duration. diaminobenzidine (DAB) solution were used to stain the sections. The staining reactions were observed carefully under a microscope and stopped with tap water in time. Finally, the sections were counterstained with hematoxylin. Negative controls were obtained by omission of the primary antibodies in all procedures of IHC.

The sections were observed by two independent pathologists. Then the staining of immunohistochemistry was classified according to the percentage of cells with a positive score for staining. Briefly, scoring method was shown below: The first, the staining intensity was divided into four levels and scored (0, negative; 1, weak; 2, moderate; 3, high). The second, the percentage of positive cells was divided into four levels and scored (0, 0% positive cells; 1,<30% positive cells; 2, 30%-70% positive cells; 3, >70% positive cells). The third, the sum of two scores for staining intensity and percentage of cells was a final score of each sample. Samples were classified as: (-) negative, the final scores were 0; (+) weak positive, the final scores were 1-2; (++) moderate positive, the final scores were 3-4; (+++) strong positive, the final scores were 5-6. HOX A5 demonstrated a membrane and cytoplasm staining. No signal was seen with negative controls.

### Western blot analysis

Cells were treated in lysis buffer (50 mM Tris-HCl, 0.5% sodium deoxycholate, 25 mM EDTA) containing protease inhibitor cocktail (Roche) for 30 min on ice. Equal amounts of protein were separated by 10% SDS-PAGE gels, and transferred onto polyvinylidene difluoride membranes. After blocked with 5% skim milk solution for 2 hours, incubated with antibodies against ADAM17(Abcam) or HOXA5(Santa Cruz) or β-actin (proteintech) at 4°C over night. The appropriate horseradish-conjugated secondary antibody was incubated for 1 hour at room temperature, the signal of protein was revealed using the chemiluminescence method (ECL, AURAGENE). Band intensities were quantified using BIO-RAD image analysis software.

### Luciferase activity assay

A 500bp sequence containing the predicted miR-224 binding site at the 3’ UTRs of ADAM17 or a 500bp sequence containing scrambled sequence cloned into the Xho I/Not I site of psiCHECK-2 vector (Promega) to generate PsiCHECK-2-ADAM17-WT and PsiCHECK-2 -ADAM17-mut vectors, respectively. Using the same approach, we also obtained the PsiCHECK-2-HOXA5-WT and PsiCHECK-2 - HOXA5-mut vectors. The whole construction process was completed by Shanghai jima pharmaceutical technology co., LTD. The corresponding sequence information is as follows: psiCHECKTM-2-ADAM17-WT:5’-TGA AGA CTG GGAA GTGAC TT-3’, psiCHECKTM-2-ADAM17-MUT:5’-TGA AGA CTG GGA GTGATT CA-3 psiCHECKTM-2-HOXA5-WT:5’-TAC CTA TAT TCC TTG TGT AAT TAA TGC TGT TGT AGAG GTGACTTG ATG AGA CAC AAC TTG TTC GAC GT-3’ psiCHECKTM-2-HOXA5-MUT: 5’-TAC CTA TAT TCC TTG TGT AAT TAA TGC TGT TGT AGAG CACTGAAC ATG AGA CAC AAC TTG TTC GAC GT-3’ HEK-293T cells were cultured in 24-well plates, and PsiCHECK-2-ADAM17-WT, PsiCHECK-2 -ADAM17-mut, PsiCHECK-2-HOXA5-WT, PsiCHECK-2 - HOXA5-mut vectors were co-transfected respectively with miR-224 mimics or negative control using Lipofectamine 2000 (Invitrogen) in 24-well plates. Luciferase assays were performed using a luciferase assay kit (Promega) according to the manufacturer’s instructions. Firefly luciferase was used for normalization.

### Cell invasion and migration assay

Transwell chamber (8μM pore diameter, Corning Costar) was used to perform the assay. The filter was coated with matrigel (BD Biosciences) for examination of cell invasion. However, the filter without coated matrigel was used for the migration assay. The trasfected cells were resuspended in serum-free medium and added 200μL cell resuspension solution (cell invasion: 1×10^5^ cells per well, cell migration: 1×10^6^ cells per well) to the hydrated transwell chambers, simultaneously, 500μL complete medium with 10% FBS was added to the bottom chambers. Cultured in the incubator for 12h (cell migration) or 24h (cell invasion), removed the cells on the upper surface of the membrane with cotton swab. Filters were fixed in 4% formaldehyde for 30 min and stained with 0.1% crystal violet for 20 min. Cells were counted under microscope. Experiments were repeated at least three times.

### Cell proliferation assay

BEL-7402 cells (4000 cells/well) were plated in 96-well plates and incubated for 24 h. MTT assay was performed to measure cell viability at 24h, 48h and 72h after transfection. The absorbance at 490 nm was measured using a microplate reader (Bio-Tek Instruments, Winooski, VT).

### Cell Cycle Analysis

BEL-7402 cells were trypsinized at 48h after transfection, and washed in PBS, fixed with 500μl of 70% ethanol at –20°C over night. Subsequently, cells were stained with propidium iodide (PI) solution at 37°C for 30 min. Analysis was performed on a FACS flow cytometer (BD Biosciences).

### Statistical analysis

The two-tailed student’s t-test, one-way ANOVA, Mann-Whitney U or Kruskal-Wallis Test were used for data analysis. The statistical analyses were done using SPSS 19.0 for windows. P<0.05 was considered significant.

## Results

### The expression of miR-224 in hepatocellular carcinoma and adjacent tissues

Expression of miR-224 was detected in 19 pairs of human hepatocellular carcinoma and adjacent liver tissues by real-time quantitative RT-PCR. Expression of miR-224 was upregulated in 2 cases of hepatocellular carcinoma tissues (>1.5 fold); It is considered to be virtually unchanged in 3 cases of hepatocellular carcinoma tissues, because the magnitude of increased or decreased miR-224 expression was less than 1.5 fold; the down-regulation of miR-224 was found in 14 cases of hepatocellular carcinoma tissues (>1.5 fold), the statistical analysis showed that the expression of miR-224 in hepatocellular carcinoma tissues was significantly higher than that in adjacent liver tissues (*P*<0.01, Mann-Whitney U test) (Fig 1A). These results suggest that miR-224 may play a role as an oncogene in hepatocellular carcinoma. Fluorescence quantitative PCR was used to detect ADAM17 expression in 19 hepatocellular carcinoma and paracancer tissues. The results showed that ADAM17 expression in 6 hepatocellular carcinoma tissues was down-regulated (the reduction was more than 1.5 times).ADAM17 expression in 8 hepatocellular carcinoma tissues did not change significantly (the amplitude of up-regulation in 5 cases or down-regulation in 2 cases was less than 1.5 times, and the ADAM17 expression in 1 case was not changed). ADAM17 expression was up-regulated (by more than 1.5 times) in 5 hepatocellular carcinoma tissues. Overall analysis showed that a total of 10 cases were upregulation, although the multiple of upregulation in 5 cases was relatively small, statistical analysis also showed that ADAM17 expression in HCC tissues was higher than that in adjacent tissues (P<0.01, Mann-Whitney U test) (Fig 1B) were statistically significant. Of course, we could also find that the number of cases that were down-regulated in fact actually exceeded the number of cases that were significant (the increase was more than 1.5 times). To achieve a more positive result, it was necessary to increase the number of cases in the later period for testing.These results suggest ADAM17 may act as an oncogene in hepatocellular carcinoma.

**Fig 1.**
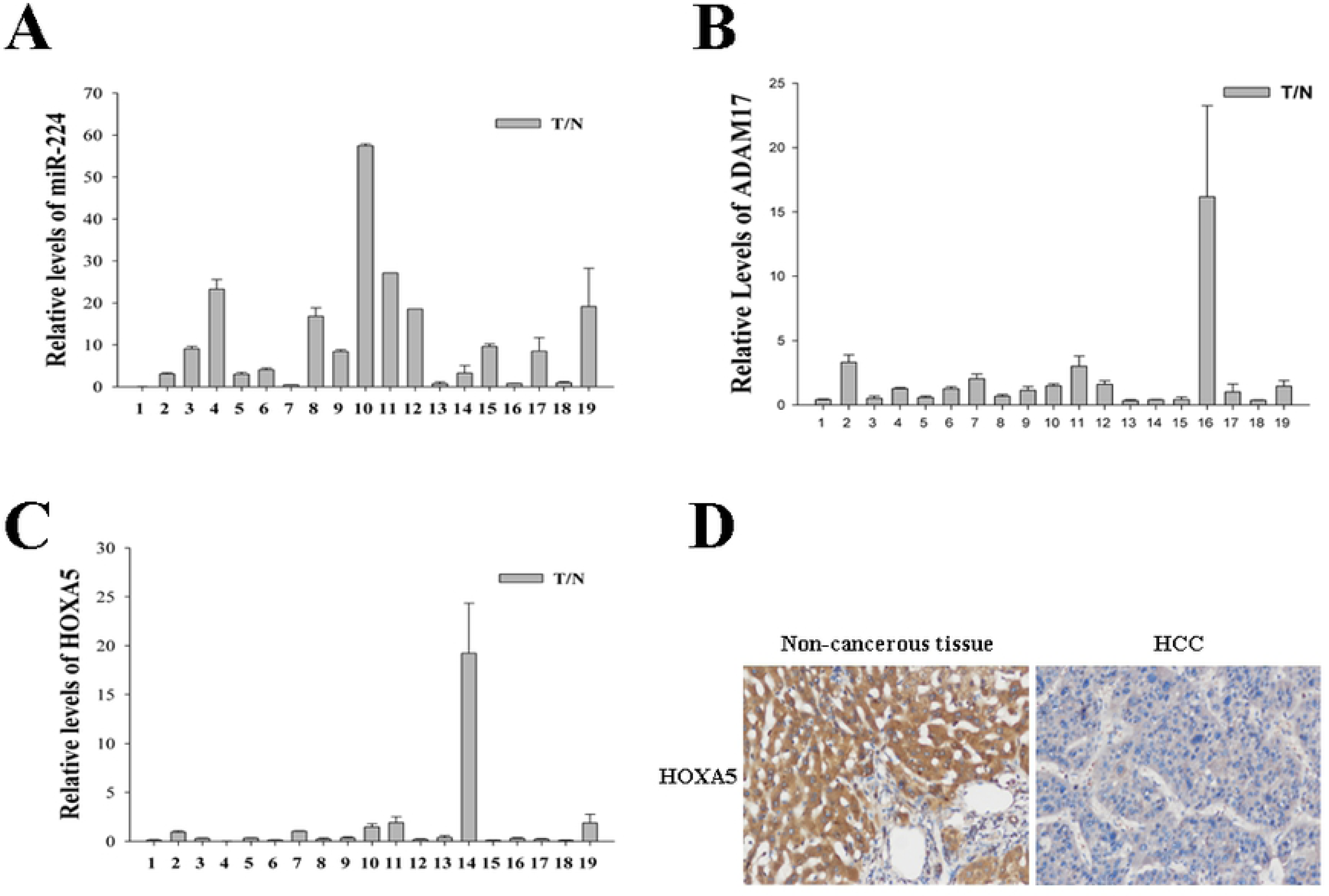
Expression of miR-224 in hepatocellular carcinoma. The relative levels of miR-224 expression in hepatocellular carcinoma tissues and adjacent liver tissues were analyzed by real time PCR, and normalized to small nuclear RNA U6.(P<0.01) (T/N-Tumor/Normal)

### Expression of miR-224 in hepatocyte cancer cell lines

Fluorescence quantitative PCR was used to detect the expression of miR-224 in SMMC-7721, BEL-7402, Huh-7 and HL-7702 cells, and the results showed significant differences in the expression of miR-224 in these cells (*P*=0.001, Kruskal-Wallis H test) (Fig S1-A). Compared with HL-7702 cells, the expression of miR-224 was up-regulated in SMMC-7721 cells (*P*=0.025, Mann-Whitney U test), while there was no difference in BEL-7402 cells (*P*=0.054, Mann-Whitney U test), the expression of miR-224 In Huh-7 cells was down-regulated (*P*=0.004, Mann-Whitney U test). The expression of miR-224 in SMMC-7721 and BEL-7402 was higher than that in Huh-7 cells (*P*=0.004, both, Mann-Whitney U test), there was no significant difference between SMMC-7721 and BEL-7402 cells (*P*=0.423, Mann-Whitney U test).

In order to understand the effect of mimics and inhibitors of miR-224 on hepatocyte cancer cells, miR-224 mimics, mimics NC, miR-224 inhibitor and inhibitor NC were transfected into SMMC-7721, BEL-7402 and Huh-7, and the expression of miR-224 was detected by fluorescence quantitative PCR 48h later. The results showed that the expression of miR-224 in SMMC-7721, BEL-7402 and Huh-7 was approximately 50, 77 and 7545 times higher than that in the control group, respectively, after transfection with miR-224 mimics.(*P*<0.05, all, Mann-Whitney U test)(Fig S1 B, C and D). While compared with the control group, the expression of miR-224 in SMMC-7721, BEL-7402 and Huh-7 decreased by approximately 19, 32 and 64 times, respectively, after transfection with miR-224 inhibitor. (*P*<0.05, all, Mann-Whitney U test)(Fig S1 B, C, D). The results suggested that the mimics and inhibitors of miR-224 could effectively promote or inhibit the expression of miR-224 in hepatocyte cancer cells.

### Identification and validation of target genes of miR-224

In order to better understand the mechanism of miR-224 in the development of HCC, TargetScan[34], miRanda[35], PITA[36] and other computational methods were used to predict the target genes of miR-224. Among the many predicted target genes, ADAM17 and HOXA5 were selected as the targets for further study. The 3’UTR region of the mRNA encoded by ADAM17 and HOXA5 genes contains sequences that are partially complementary to miR-224 (Fig 2).

**Fig 2.**
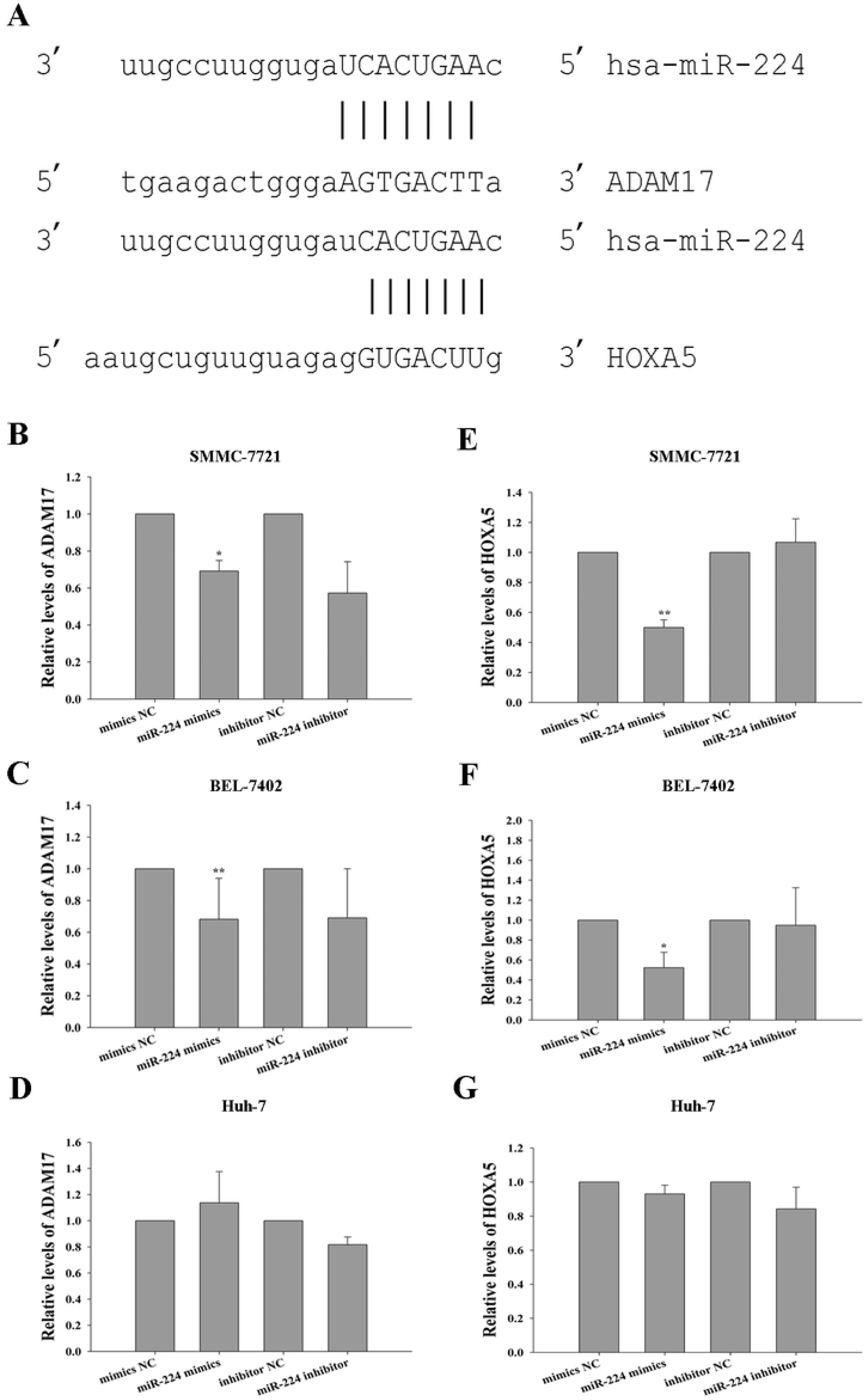
The predicted binding sites of miR-224 and target genes ADAM17 and HOXA5. The binding site of ADAM17 to miR-224 was predicted by miRanda (V1.9), and the binding site of HOXA5 to miR-224 was predicted by TargetScan (v4.1).

miR-224 mimics, mimics NC, miR-224 inhibitor and inhibitor NC were transfected into SMMC-7721, BEL-7402 and Huh-7 cells, respectively, and the mRNA expression levels of ADAM17 and HOXA5 were detected by fluorescence quantitative PCR and Western blottig 48h later. The results showed that the overexpression of miR-224 in SMMC-7721 and BEL-7402 could significantly reduce the mRNA expression of ADAM17 and HOXA5 (*P*<0.05, all, the two-tailed Student’s t-test)(Fig 3), while the overexpression of miR-224 in Huh-7 had no significant effect on the mRNA expression of ADAM17 and HOXA5 (*P*>0.05, all,The two-tailed Student’s t-test (Fig 3). Inhibition of miR-224 expression in three hepatocyte cancer cells could not effectively increase the mRNA expression levels of ADAM17 and HOXA5 (*P*>0.05, all, the two-tailed Student’s t-test)(Fig 3).

**Fig 3.**
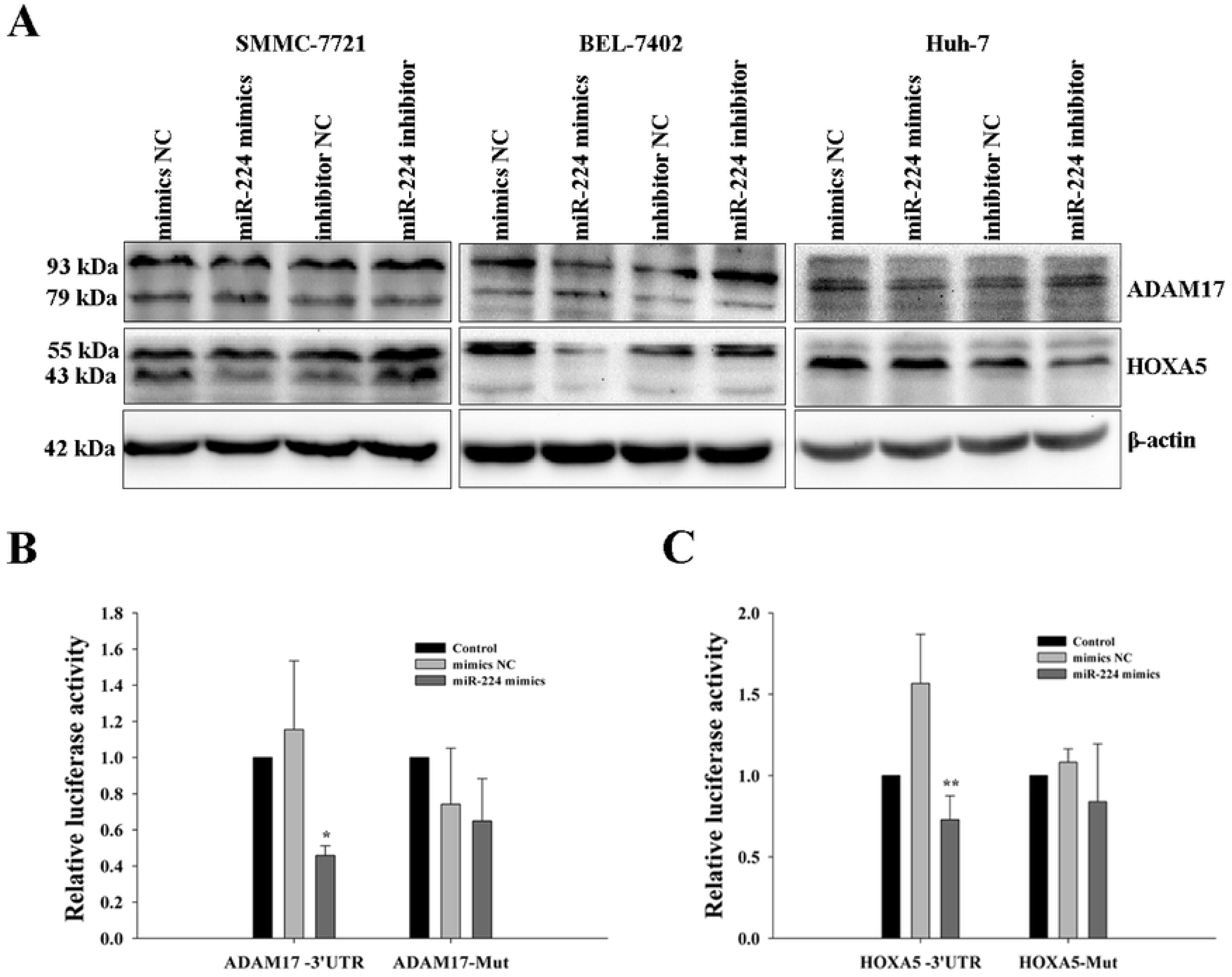
MRNA expression changes of target genes ADAM17 and HOXA5 after overexpression or inhibition of miR-224. A,B,C. MRNA expression changes of ADAM17 in SMMC-7721, BEL-7402 and Huh-7 cells after overexpression or inhibition of miR-224. D,E,F. MRNA expression changes of HOXA5 in SMMC-7721, BEL-7402 and Huh-7 cells after overexpression or inhibition of miR-224. (*,P< 0.05, **,P< 0.01)

The overexpression of miR-224 could cause the protein expression level of ADAM17 in SMMC-7721, BEL-7402 and Huh-7 cells to decline, the protein expression level of HOXA5 in SMMC-7721 and BEL-7402 cells decreased, but it had no significant effect on the protein expression level of HOXA5 in Huh-7 cells. The inhibition of miR-224 could cause the expression level of ADAM17 in SMMC-7721, BEL-7402 and Huh-7 cells to rise, the expression level of HOXA5 in SMMC-7721 and BEL-7402 cells to rise, but had no significant effect on the expression level of HOXA5 in Huh-7 cells (Fig 4).

**Fig 4.**
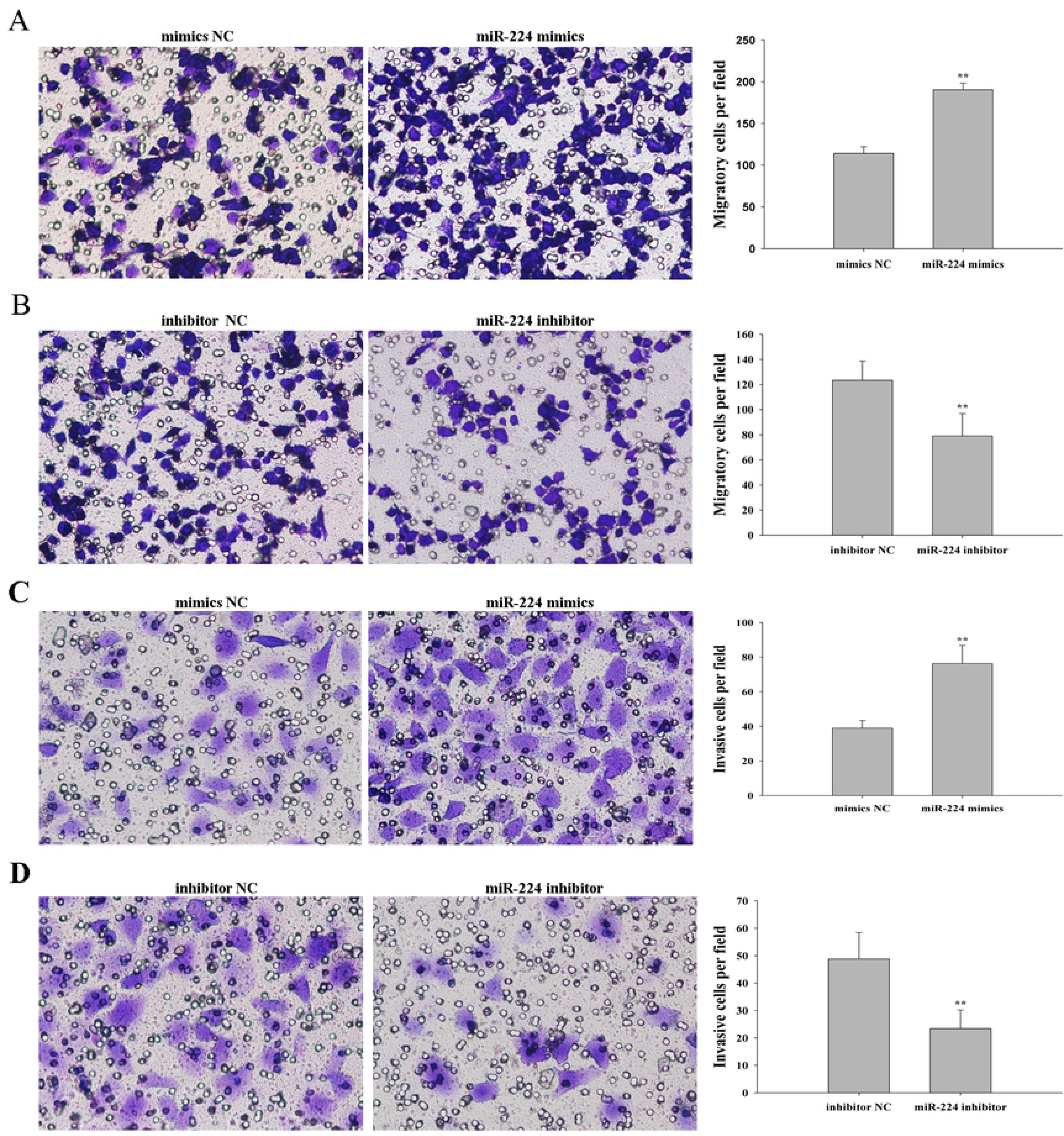
Effect of overexpression or inhibition of miR-224 on protein expression levels of its target genes ADAM17 and HOXA5. SMMC-7721, BEL-7402 and HuH-7 cells were transfected with miR-224 mimics, mimics negative control(NC), miR-224 inhibitor and inhibitor negative control(NC), at 48 hours after trasfecton, protein expression of ADAM17 and HOXA5 were analyzed by Western blotting, as compared to negative control (NC). ADAM17 protein has two subunits, the molecular weight of 93 kDa and 79 kDa, respectively. HOXA5 protein has also two subunits, the molecular weight of 55 kDa and 43 kDa, respectively.

In order to detect the direct interaction between miR-224 and its target genes ADAM17 and HOXA5, we co-transfected miR-224 mimics or mimics NC with wild-type or mutant vectors of ADAM17 and HOXA5 into 293T cells, and then detected the changes of double luciferase of the vector. We found that the fluorescence activity of the wild-type reporting genomes of ADAM17 and HOXA5 was significantly lower than that of the control group (*P*<0.05, one-way ANOVA) (Fig 5). This result suggests that miR-224 can directly regulate ADAM17 and HOXA5.

**Fig 5.**
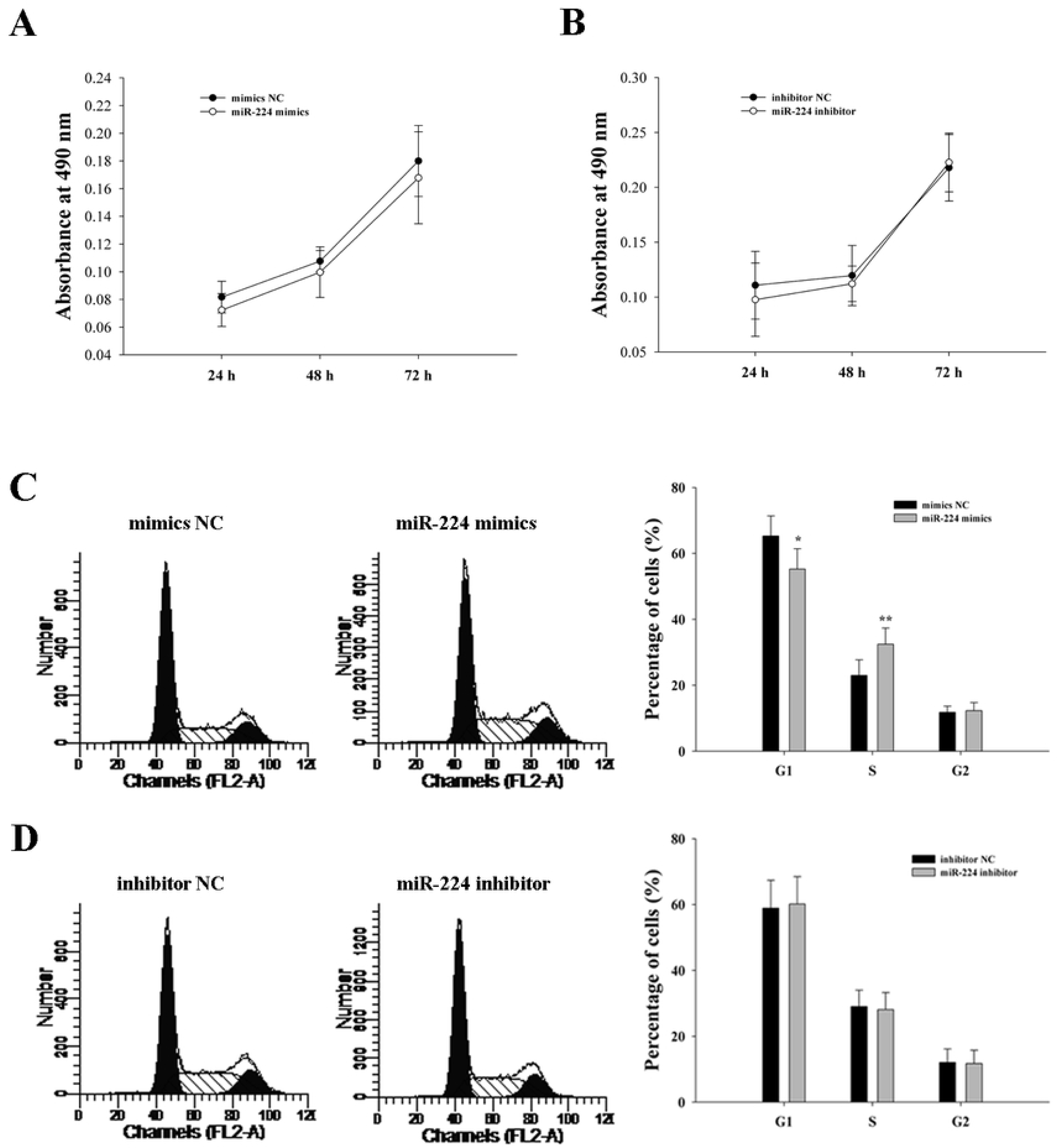
Double luciferase activity of target genes ADAM17 and HOXA5 of miR-224 was detected. A. Luciferase activity of wild type or mutant ADAM17 3’UTR reporter gene in HEK-293T cells transfected with miR-224 mimics or mimics negative control(NC) was detemined. The results showed that luciferase activity of wild type ADAM17 3’UTR transfected with miR-224 mimics was decreased compare to others group in HEK-293T cells. B. Luciferase activity of wild type or mutant HOXA5 3’UTR reporter gene in HEK-293T cells transfected with miR-224 mimics or mimics negative control(NC) was detemined. The results showed that luciferase activity of wild type HOXA5 3’UTR transfected with miR-224 mimics was decreased compare to others group in HEK-293T cells. (*, P<0.05, **, P<0.01)

### Regulation of miR-224 to Cell Migration and Invasion Viability

Combining the effects of miR-224 mimics and inhibitor on three strains of HCC, the regulatory effects of two target genes, and the biological characteristics of cancer cells, BEL-7402 cells were selected to complete the biological function test of miR-224 overexpression or inhibition.

We evaluated the effect of miR-224 on cell migration and invasion ability in vitro by enhancing or silencing the expression of miR-224. 48h after transfection with miR-224 mimics or mimic NC into BEL-7402 cells, the effect of miR-224 overexpression on the migration and invasion ability of BEL-7402 cells was detected by Transwell assay. The results showed that compared with the mimics NC group, the migration (*P*<0.01, the two-tailed Student’s t-test) (Fig 6A) and invasion (*P*<0.01, the two-tailed Student’s t-test) (Fig 7A) capabilities of the miR-224 mimics group were significantly enhanced. 48h after transfection of BEL-7402 cells with miR-224 inhibitor or inhibitor NC, the effect of miR-224 silencing on the migration and invasion ability of bel-7402 cells was determined by Transwell assay. The results showed that compared with the inhibitor NC group, the ability of miR-224 inhibitor group to migrate (P <0.01, the two-tailed Student’s t-test) (Fig 6B) and to invade (P <0.01, the two-tailed Student’s t-test) (Fig 7B) was significantly reduced.

**Fig 6.**
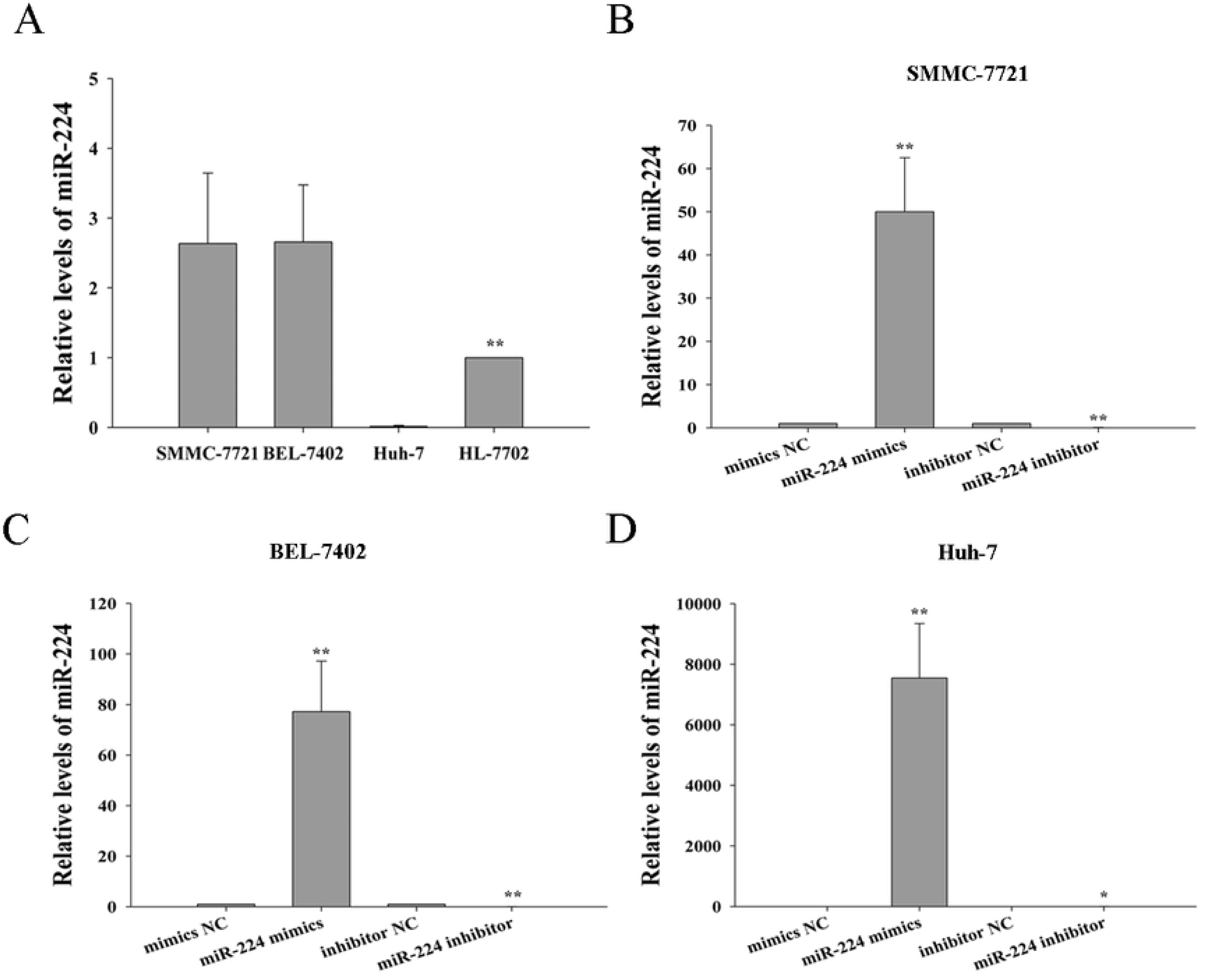
miR-224 affects the migration ability of BEL -7402 cells. A. The overexpression of miR-224 promoted the migration ability of BEL-7402 cells. B. miR-224 silencing inhibited the migration of BEL-7402 cells. (*,P< 0.05, **,P< 0.01)

**Fig 7.** miR-224 affects the invasion ability of BEL -7402 cells. A. The overexpression of miR-224 promoted the invasion ability of BEL-7402 cells. B. miR-224 silencing inhibited the invasion of BEL-7402 cells. (*,P< 0.05, **,P< 0.01)

### Effect of miR-224 on proliferation of BEL-7402 cells

In order to detect whether the abnormal expression of miR-224 had an impact on the growth of cancer cells during the development of HCC, we transfected miR-224 mimics, mimic NC, miR-224 inhibitor and inhibitor NC into BEL-7402 cells, and detected the cell activity by MTT method at 24h, 48h and 72h after transfection, respectively. The results showed that overexpression or silencing of miR-224 had no significant effect on the proliferation of BEL-7402 cells (*P*>0.05, the two-tailed Student’s t-test)(Fig 8).

**Fig 8.** Effect of miR-224 on proliferation of bel-7402 cells. A. The overexpression of miR-224 had no effect on the proliferation of BEL-7402 cells. B. The silencing of miR-224 had no effect on the proliferation of BEL-7402 cells.

### Effect of miR-224 on the cell cycle of BEL-7402

In order to detect whether the abnormal expression of miR-224 had an impact on the distribution of cell cycle during the development of HCC, we transfected miR-224 mimics, mimic NC, miR-224 inhibitor and inhibitor NC into BEL-7402 cells, and detected the cell cycle changes by flow cytometry 48h after transfection, we found that miR-224 mimics could reduce the cell percentage in G1 phase of the cell cycle (*P* <0.05, the two-tailed Student’s t-test) (Fig 9A) and increase the cell percentage in S phase (*P*<0.01, the two-tailed Student’s t-test) (Fig 9A), but had no significant effect on the cell percentage composition in G2 phase (*P*=0.682,The two-tailed Student’s t-test (Fig 9A).In addition, after miR-224 was silenced, flow results showed that the cell percentage of miR-224 inhibitor group in G1, S and G2 phases did not change significantly compared with that of inhibitor NC group (*P*>0.05, all, the two-tailed Student’s t-test)(Fig 9B).These results suggest that miR-224 overexpression can promote the proliferation of bel-7402 cells.

**Fig 9.** The effect of miR-224 on the cell cycle of bel-7402. A. The effect of overexpression of miR-224 on the cell cycle of BEL-7402, Cell cycle analysis was performed by FCM (flow cytometry), results showed that miR-224 mimics could reduce the cell percentage in G1 phase of the cell cycle and increase the cell percentage in S phase, but had no significant effect on the cell percentage composition in G2 phase B. The effect of miR-224 silencing on the cell cycle of BEL-7402, Cell cycle analysis was performed by FCM (flow cytometry), results showed that the cell percentage of miR-224 inhibitor group in G1, S and G2 phases did not change significantly compared with that of inhibitor NC group. (*.P< 0.05, **P< 0.01).

## Discussion

Hepatocellular carcinoma (HCC) is one of the most common malignant tumors,Its morbidity and mortality rates have been on the rise in recent years[37], the high recurrence or metastasis rate after surgery may hinder the improvement of the survival time of hepatocellular carcinoma patients[37]?In order to get a better method to detect hepatocellular carcinoma, a biological target is needed to effectively screen high-risk patients for early diagnosis so that interventions can be made earlier, increasing the likelihood of a cure.MiRNAs appear as important regulators of genes, and ongoing studies on their functions and target genes reveal that MiRNAs play a key role in several important cellular functions. MiRNAs possess several characteristics that make them possible targets for early diagnosis of tumors or powerful tools for gene therapy, which will bring new hope for the diagnosis and treatment of tumors[38], this is very attractive to researchers.

miR-224 is present in mammals, including humans, and is located on the X chromosome[39]. It has been reported that miR-224 is abnormally expressed in a variety of tumors, suggesting that it may play an important role in the biological process of tumors.According to the different malignant phenotypes of tumor cells, the biological role of miR-224 is also different, and it also shows contradictory properties in different studies. Several reports have confirmed that the expression of miR-224 in hepatocellular carcinoma is up-regulated, and overexpression of miR-224 can promote proliferation and invasion of hepatocellular carcinoma cells[30, 40] .However, in prostate cancer, the expression of miR-224 is down-regulated and inhibits the migration and invasion of prostate cancer cells by regulating TPD52[41]. In addition, overexpression of miR-224 in adult neurohemangiomas can inhibit cell proliferation[42]. These results suggest that the mechanism of miR-224 in tumors is complex, requiring further study.

In this study, we used a variety of methods to identify and confirm the target genes of miR-224 and their mechanism of action in hepatocarcinoma. The up-regulation of miR-224 can reduce the mRNA and protein expression levels of ADAM17 and HOXA5, and the double luciferase experiment further proved that miR-224 has a direct targeting regulation effect on its target genes ADAM17 and HOXA5. Importantly, in the functional test of miR-224, we found that overexpression of miR-224 significantly enhanced the migration and invasion ability of Bel-7402 cells.In contrast, silencing of miR-224 inhibited the migration and invasion of Bel-7402 cells. In addition, overexpression of miR-224 can change the cell cycle of Bel-7402 cells, and the cell proportion of S phase increases significantly, while the cell proportion of G1 phase decreases significantly, while the cell proportion of G2 phase remains unchanged, suggesting that overexpression of miR-224 can promote the proliferation of Bel-7402 cells.The reason for this result may be that overexpression of miR-224 causes changes in genes related to GI/S transformation and DNA replication, but further studies are needed to prove this.Paradoxically, the silencing of miR-224 had no meaningful effect on the cell cycle. In addition, in the analysis of MTT experimental results, we failed to find evidence of the effect of overexpression or silencing of miR-224 on the proliferation of Bel-7402 cells. This may be due to the lack of an experimental strategy for stable overexpression or silencing of genes that hinders the analysis of these experiments, although individual differences between cell models should also be fully considered.

Using bioinformatics to predict miRNAs target genes is a convenient method for preliminary screening of target genes, which can provide us with many candidate target genes. Among these predicted target genes, many target genes have been experimentally proved to play very important roles in the occurrence and development of tumors. These target genes of miR-224 with rich functional categories further confirmed that miR-224 may be directly or indirectly involved in the biological process of tumors. It remains a difficult task to fully understand the mechanisms by which miRNAs dysfunction may promote tumours, and although bioinformatics tools can help reveal the putative mRNA target genes, experimental data confirmation remains a necessary tool. Some target genes of miR-224 have been reported, including RKIP[43], p21[44], Cdc42[45], TPD52[41], TRIB1[46]?HOXD10[47], SMAD4[48], API-5[31] and CXCR4[49]. Some of these target genes promote tumor development, while others do not. In this study, the data showed that miR-224 was involved in the migration and invasion of liver cancer cells, and its direct regulatory relationship with the two target genes ADAM17 and HOXA5 has also been proved. We need to further analyze which pathway actually plays a regulatory role in the migration and invasion of liver cancer cells. Our previous studies have found that ADAM17 is overexpressed in HCC tissues, and this abnormal expression is associated with microvascular invasion of HCC[50]. The ANG II-EGFR signaling pathway mediated by ADAMs (ADAM9 and ADAM17) can accelerate the proliferation and infiltration of hepatocyte cancer cells huh-7 and PLC/PRF/5, and ADAMs inhibitors can inhibit the role of this pathway[51].In this study, upregulation of miR-224 inhibited the expression of ADAM17, while the decreased expression of ADAM17 inhibited the migration and invasion of cancer cells.Therefore, the enhancement of migration and invasion of Bel-7402 cells by miR-224 may not be realized by directly regulating the target gene ADAM17.

HOXA5 is down-regulated in hepatocellular carcinoma, suggesting that it may be a tumor suppressor gene in hepatocellular carcinoma.In addition, it has also been reported that HOXA5 is also low-expressed in non-small cell lung cancer [52]and oral squamous cell carcinoma[53]. As the upstream regulatory network molecule of AMELY (Amelogenin y-linked), HOXA5 is involved in the regulation of cell proliferation and angiogenesis and other biological functions of hepatocellular carcinoma[54]. Inhibition of HOXA5 can lead to increased proliferation, migration and invasion of lung cancer A549 cells[55]. In this study, overexpression of miR-224 reduced the expression levels of HOXA5 mRNA and protein, while silencing of miR-224 increased the expression levels of HOXA5 mRNA and protein. This is consistent with the trend that miR-224 regulates the migration and invasion of hepatocyte cancer cells BEL-7402. Therefore, miR-224 may enhance the migration and invasion of cancer cells by directly regulating the expression of target gene HOXA5.

Tumor invasion involves several different steps[56], miR-224 may also regulate other target genes associated with tumor invasion, which is certainly a very complex information network.Our findings indicate that miR-224 plays a very important role in the occurrence and development of hepatocellular carcinoma. Further understanding of the mechanism of action of miR-224 and other target genes in hepatocellular carcinoma is conducive to the exploration of its exact role in the initiation and progression of hepatocellular carcinoma, which provides us with new research directions and challenges.

## Acknowledgements

We thank Teacher Xiaojing Yang and Dr Zhaojun Duan for her excellent technical assistance.

## Supporting information

**S1 Fig Expression of miR-224 in hepatocyte cancer cells**. A. Expression of miR-224 in hepatocyte cancer cells SMMC-7721, BEL-7402, Huh-7 and normal human hepatocytes HL-7702. B,C,D. after transfection of miR-224 mimics and inhibitor into SMMC-7721, BEL-7402 and Huh-7, the expression level of miR-224 was significantly increased or decreased compared with the negative control group.(*,P< 0.05, **,P< 0.01)

